# Impact of an Oxidative RNA Lesion on in vitro Replication Catalyzed by SARS-CoV-2 RNA-dependent RNA Polymerase

**DOI:** 10.1101/2024.10.13.618053

**Authors:** Masataka Akagawa, Kaoru Sugasawa, Kiyoe Ura, Akira Sassa

## Abstract

The production of reactive oxygen species in response to RNA virus infection results in the oxidation of viral genomic RNA within infected cells. These oxidative RNA lesions undergo replication catalyzed by the viral replisome. G to U transversion mutations are frequently observed in the SARS-CoV-2 genome and may be linked to the replication process catalyzed by RNA-dependent RNA polymerase (RdRp) past the oxidative RNA lesion 7,8-dihydro-8-oxo-riboguanosine (8-oxo-rG). To better understand the mechanism of viral RNA mutagenesis, it is crucial to elucidate the role of RdRp in replicating across oxidative lesions. In this study, we investigated the RNA synthesis catalyzed by the reconstituted SARS-CoV-2 RdRp past a single 8-oxo-rG. The RdRp-mediated primer extension was significantly inhibited by 8-oxo-rG on the template RNA. During the blockage of the extension reaction, the rate of product release by RdRp was extremely slow, indicating that the RdRp machinery remained bound to the replicating primer/template RNA. Once RdRp was able to bypass 8-oxo-rG, it preferentially incorporated rCMP, with a lesser amount of rAMP opposite 8-oxo-rG. In contrast, RdRp demonstrated greater activity in extending from the mutagenic rA:8-oxo-rG terminus compared to the lower efficiency of extension from the rC:8-oxo-rG pair. Based on steady-state kinetic analyses for the incorporation of rNMPs opposite 8-oxo-rG and chain extension from rC:8-oxo-rG or rA:8-oxo-rG, the relative bypass frequency for rA:8-oxo-rG was found to be seven-fold higher than that for rC:8-oxo-rG. Therefore, the properties of RdRp indicated in this study may contribute to the mechanism of mutagenesis of the SARS-CoV-2 genome.

## Introduction

RNA viruses are characterized by extremely high mutation rates, ranging from 10^−4^ to 10^−6^ substitutions per nucleotide per cell infection (s/n/c), which is one million times higher than that of eukaryotic host cells(1, 2). Such elevated mutation rates ensure the evolvability of these viruses (2). For example, the mutation rate of influenza A viruses (IVA), which possess a single-stranded RNA genome, is approximately1.5 × 10^−5^ (s/n/c) (3). Similarly, the mutation rate for the human immunodeficiency virus (HIV) genome ranges from 10^−4^ to 10^−5^ (s/n/c)(4). In the case of the Hepatitis C virus (HCV), it is estimated to be between 1.6 and 6.2 × 10^−5^ per nucleotide per genome replication (5). Compared to other RNA viruses, coronaviruses (CoVs), which cause respiratory diseases in humans (6), exhibit lower mutation rates due to the presence of 3′-exoribonuclease in the replication complex (7). Over the past 25 years, highly pathogenic human coronaviruses have emerged, including the severe acute respiratory syndrome coronavirus (SARS-CoV, 2002), Middle East respiratory syndrome coronavirus (8), and severe acute respiratory syndrome coronavirus 2 (SARS-CoV-2, 2019). As of today, more than 775 million patients have been infected with SARS-CoV-2 (9). The spontaneous mutation rate of SARS-CoV-2 is estimated to be approximately 1.3 × 10^−6^, which is similar to that of other betacoronavirus (10, 11).

Among the known mutations in ssRNA viruses, A to G transition mutations are frequently observed and are likely induced by the host RNA editing enzyme ADAR1 (12). ADAR1 exhibits hydrolytic deamination activity, converting adenosine to inosine in dsRNA. Since inosine is recognized as guanosine during replication, A-to-I modifications lead to an increased frequency of A to G mutations in RNA genomes. In the SARS-CoV-2 genome, C to U transition mutations are also frequently observed (13), and these mutations have been reported to result from deamination by the host cytidine deaminases APOBEC1, APOBEC3A, and APOBEC3G (14, 15). The expression of these APBECs has been shown to enhance the replication of the SARS-CoV-2 genome. Interestingly, G to U transversion mutations are the second most frequently observed mutations in the SARS-CoV-2 genome (13, 16–19). These mutations are believed to be associated with the mutagenic effects of reactive oxygen species (ROS), which generate the oxidative base lesion 7,8-dihydro-8-oxoguanine (8-oxoG) (20, 21).

Upon infection, RNA viruses release their genomic RNA into the cytoplasm of host cells (22, 23). This viral infection triggers the generation of ROS through mitochondrial respiratory reactions (24, 25). Additionally, infection-associated ROS are produced by NADPH oxidase, which is activated by Toll-like receptor 7 in response to viral single-stranded RNA (ssRNA) (25–27). Moreover, SARS-CoV-2 infection exacerbates the expression of proinflammatory cytokines such as IL-1, IL-6, and IL-18, leading to an overproduction of ROS (28). Excessive levels of ROS can damage various cellular components, including proteins, lipids, DNA, and RNA, causing organ dysfunction and contributing to disease progression (29). Therefore, the viral RNA genome also becomes susceptible to oxidative damage from endogenous cellular processes during infection. Among the nucleic acid bases found in DNA and RNA, guanine has the lowest oxidative potential compared to other bases (30), with 8-oxoG recognized as the most prevalent form of oxidative base damage (31, 32). In the genomic DNA of both prokaryotes and eukaryotes, 7,8-dihydro-8-oxo-2’-deoxyguanosine (8-oxo-dG) exhibits dual coding potential due to its *anti*- and *syn*-conformations, enabling it to form Watson–Crick pairs with cytosine and Hoogsteen pairs with adenine, respectively (33). If the 8-oxo-dG:dA mispair is left unrepaired, a dT:dA pair is generated during the second round of replication, resulting in a G:C to T:A transversion mutation. To mitigate the harmful effects of oxidative DNA lesions, mammalian cells have evolved robust DNA repair mechanisms, including OGG1, NEIL1, NTH1, and MYH (34, 35). Additionally, both prokaryotes and eukaryotes possess various DNA polymerases capable of bypassing different DNA lesions during replication (36–41). In contrast, the viral RNA genome is synthesized by a single RNA polymerase, meaning that the oxidized RNA base, 7,8-dihydro-8-oxoriboguanosine (8-oxo-rG), is replicated solely by the viral replisome. The effects of RNA lesions on viral RNA replication remains poorly understood.

RNA-dependent RNA polymerase (RdRp) is responsible for replicating the RNA viral genome (42). The SARS-CoV-2 RdRp is a multi-subunit complex composed of three different nonstructural proteins (nsps): nsp7, nsp8, and nsp12. The catalytic subunit, nsp12, features a C-terminal polymerase domain and an N-terminal nidovirus RdRp-associated nucleotidyltransferase (NiRAN) domain (43, 44). The polymerase domain resembles a right hand, consisting of the thumb, fingers, and palm subdomains (43, 44). While nsp12 possesses only minimal polymerase activity on its own, the addition of the accessory subunits nsp7 and nsp8 significantly enhances its catalytic activity (45–48). Several studies have reported the molecular basis of RNA synthesis and nucleotide selectivity by SARS-CoV-2 RdRp (49–53). Additionally, a recent study revealed the activity of RdRp on various natural RNA modifications (54). However, little is known about RdRp’s behavior in response to RNA lesions during replication. In this study, we investigated the RNA synthesis catalyzed by the reconstituted SARS-CoV-2 RdRp past a single 8-oxo-rG on the template RNA. Our observations showed that 8-oxo-rG strongly impeded RdRp-mediated primer extension. During this retardation of the extension reaction, the rate of product release by RdRp was extremely slow, indicating that the RdRp machinery remained bound to the replicating primer/template RNA. When RdRp was able to insert rNMP opposite 8-oxo-rG, it predominantly incorporated rCMP, with a lesser amount of rAMP opposite 8-oxo-rG. In contrast, RdRp efficiently extended from the mutagenic rA:8-oxo-rG terminus compared to the non-mutagenic rC:8-oxo-rG pair. Overall, the bypass frequency for rA:8-oxo-rG was higher than that for rC:8-oxo-rG. A possible mechanism by which RdRp contributes to mutagenesis via oxidation of the SARS-CoV-2 genome is discussed.

### Experimental procedures

#### Expression and purification of the SARS-CoV-2 RdRp complex

The expression and purification of the recombinant RdRp complex were conducted following a previously published study (55). Briefly, the plasmid pRSFDuet-1 (14 × His-TEV-nsp8/nsp7)(nsp12) was transformed into the BL21 Star (DE3) *E. coli* strain. Cells were cultured in LB medium supplemented with 100 µg/mL kanamycin at 37°C with 150 rpm agitation overnight. Subsequently, 20 mL of the overnight culture was diluted into 2L of fresh LB and further agitated at 30°C until the culture reached an absorbance of 0.5 at 595 nm. The overexpression of the nsp12/14 × His-TEV-nsp8/nsp7 subunits was induced by adding 0.05 mM isopropyl-1-thio-β-D-galactopyranoside at 16°C with 150 rpm agitation overnight. The harvested cells were then lysed in HisTrap buffer A [50 mM Na-HEPES (pH 8.0), 500 mM NaCl, 10% Glycerol, 10 mM Imidazole, 5 mM β-mercaptoethanol, 25-29 U/µL Benzonase, and complete EDTA-free Protease Inhibitor (Roche)) via sonication on ice. Soluble proteins were obtained through ultracentrifugation at 131,500 g for 30 min at 4°C. Following ultracentrifugation, the supernatant containing the nsp12/14 × His-nsp8/nsp7 proteins was loaded onto a TALON Metal Affinity Resin column (Clontech). The column was washed with eight column volumes of HisTrap buffer A, and the proteins were eluted with four column volumes of HisTrap buffer B [50 mM Na-HEPES (pH 8.0), 500 mM NaCl, 10% Glycerol, 360 mM Imidazole, 5 mM β-mercaptoethanol). The eluted fraction was diluted fivefold in a 50mM Na-HEPES (pH 8.0) solution before being loaded onto a Capto HiRes Q 5/50 anion-exchange chromatography column using an ÄKTA Go protein purification system (Cytiva). After loading the protein fraction, the column was washed with HiTrap Q buffer A [50 mM Na-HEPES (pH 8.0), 150 mM NaCl, 10% Glycerol, 5 mM β-mercaptoethanol), while the protein was eluted using a linear gradient of NaCl created by mixing HiTrap Q buffer A with HiTrap Q buffer B [50 mM Na-HEPES (pH 8.0), 1 M NaCl, 10% Glycerol, 5 mM β-mercaptoethanol). After elution, 200 µL of 2 mg/mL TEV protease was added to the combined purest fractions containing the RdRp complex, and the mixture was shaken at 4°C overnight. Subsequently, the TEV protease-treated protein was loaded onto a TALON Metal Affinity Resin column to remove the detached His-tag. The flow-through fraction was concentrated to 500 µl by ultrafiltration at 5,000 g using Amicon Ultra-4 centrifugal filter units with a 30,000 NMWL (EMD Millipore). The concentrated protein fraction was subjected to Superdex 200 10/300, equilibrated with gel filtration buffer [20 mM Na-HEPES (pH 8.0), 300 mM NaCl, 1 mM MgCl_2_, 10% Glycerol, and 5 mM β-mercaptoethanol). Fractions containing RdRp were then combined and concentrated to 360 µl by ultrafiltration at 5,000 g using Amicon Ultra-4 centrifugal filter units with a 30,000 NMWL (EMD Millipore) at 4°C. The purified recombinant RdRp was aliquoted, flash-frozen in liquid nitrogen, and stored at −80°C.

#### Preparation of the primer/template RNA

The 40-mer RNA template (5′-CUAUCCCCAUGUGAUUUUAAUAXCUUCUUAGGAGAAUGAC-3′, where X represents G or 8-oxo-rG) corresponding to the 3′ end of the SARS-CoV-2 genome, along with 5′-Cy3-labeled RNA primers, was synthesized by Tsukuba Oligo Service Co., Ltd. (Ibaraki, Japan). To prepare the primer/template RNA (P/T RNA) for the primer extension reactions, the primers were annealed to the templates at a 1:1.2 molar ratio in annealing buffer [10 mM Tris-HCl (pH 8.0), 50 mM NaCl, 1mM EDTA). The sample was then incubated for 5 min at 70°C and subsequently cooled to 25°C at a rate of 5°C/min. For primer extension past the lesion illustrated in Figure 1, the RNA template was annealed to a 14-mer 5′-Cy3-labeled primer (5′-GUCAUUCUCCUAAG-3′). For single-nucleotide incorporation opposite the lesion depicted in Figure 2, the RNA template was annealed to a 5′-Cy3-labeled 17-mer primer (5′-GUCAUUCUCCUAAGAAG-3′). For facilitate extension from matched (rC:rG or rC:8-oxo-rG) or mismatched (rA:rG or rA:8-oxo-rG) primer termini shown in Figure 3, the RNA template was annealed to a 5′-Cy3-labeled 18-mer primer (5′-GUCAUUCUCCUAAGAAGN-3′, N represents C or A).

**Fig 1.**
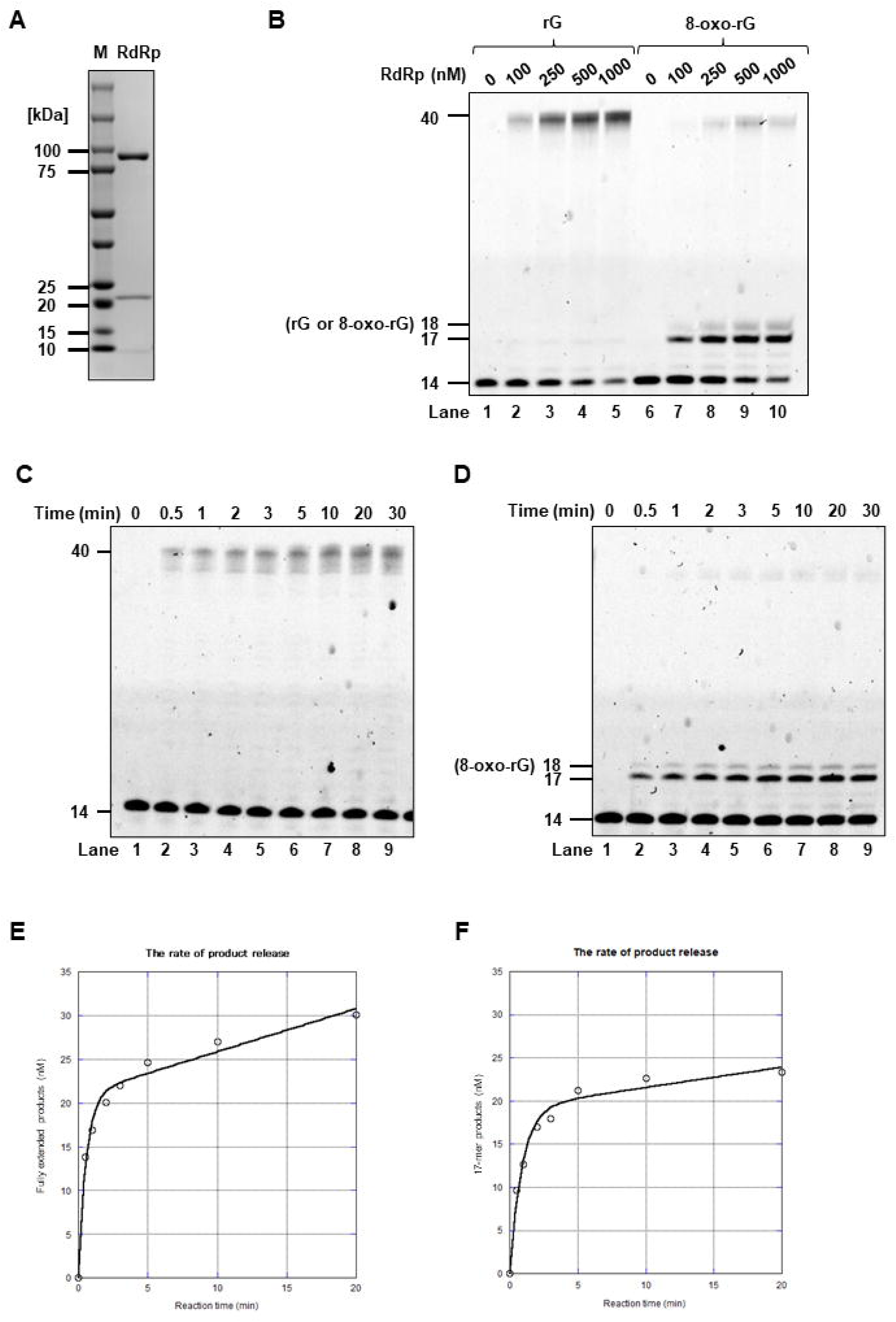
8-Oxo-rG strongly retards the RdRp-mediated primer extension. **(A)** SDS–PAGE analysis of the purified RdRp complex. The purified RdRp complex was subjected to 5%– 20% polyacrylamide gel electrophoresis and visualized using Coomassie staining. **(B)** Primer extension catalyzed by RdRp. The unmodified (lane 1–5) or 8-oxo-rG-modified (lane 6–10) 40-mer template RNA were annealed to a 5′-Cy3-labeled 14-mer primer. Reactions were catalyzed by varying concentrations of RdRp (100, 250, 500, or 1,000 nM) and conducted for 30 min in the presence of 50 µM of four rNTPs. Time-courses analysis for primer extension reactions catalyzed by RdRp used either the unmodified **(C)** or 8-oxo-rG-modified **(D)** 40-mer template annealed with a 5′-Cy3-labeled 14-mer primer. These reactions were performed with 100 nM RdRp and 100 µM of the four rNTPs for varying reaction durations, as indicated (0.5, 1, 2, 3, 5, 10, 20, 30 min). The plots of fully extended products with the unmodified template/primer **(E)** or the 17-mer products with the 8-oxo-rG-modified template/primer **(F)** were fitted to Equation 1, as described in the Experimental procedures section. The resulting parameters are tabulated in Table 1.

**Fig 2.**
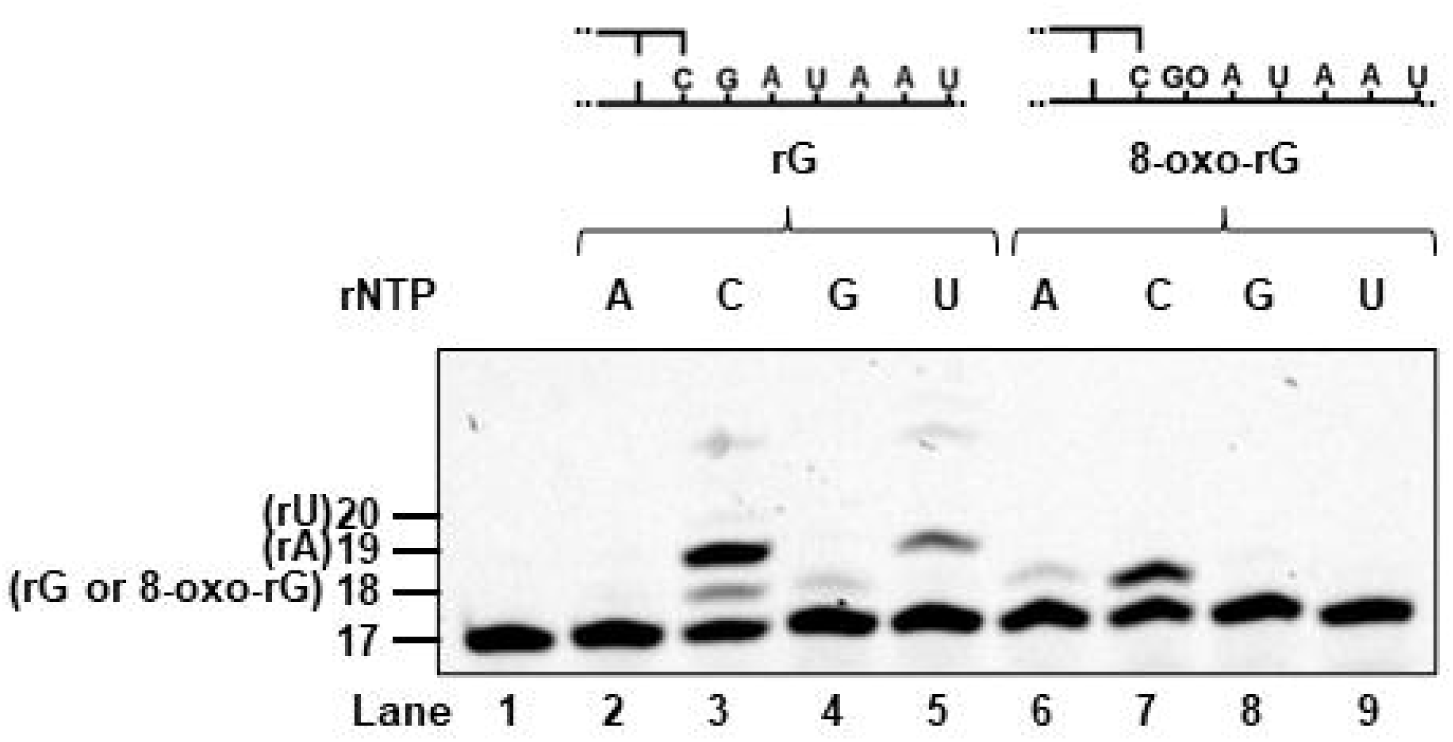
Incorporation of rNMPs opposite 8-oxo-rG by RdRp. Single-nucleotide incorporation catalyzed by RdRp was analyzed using unmodified (lane 2-5) or 8-oxo-rG (GO)-modified (lane 6–9) 40-mer template RNA, which was annealed to a 5′-Cy3-labeled 17-mer primer. Reactions were conducted with 500 nM RdRp for 30 min in the presence of 50 µM of a single rNTP (rATP, rCTP, rGTP, or rUTP).

**Fig 3.**
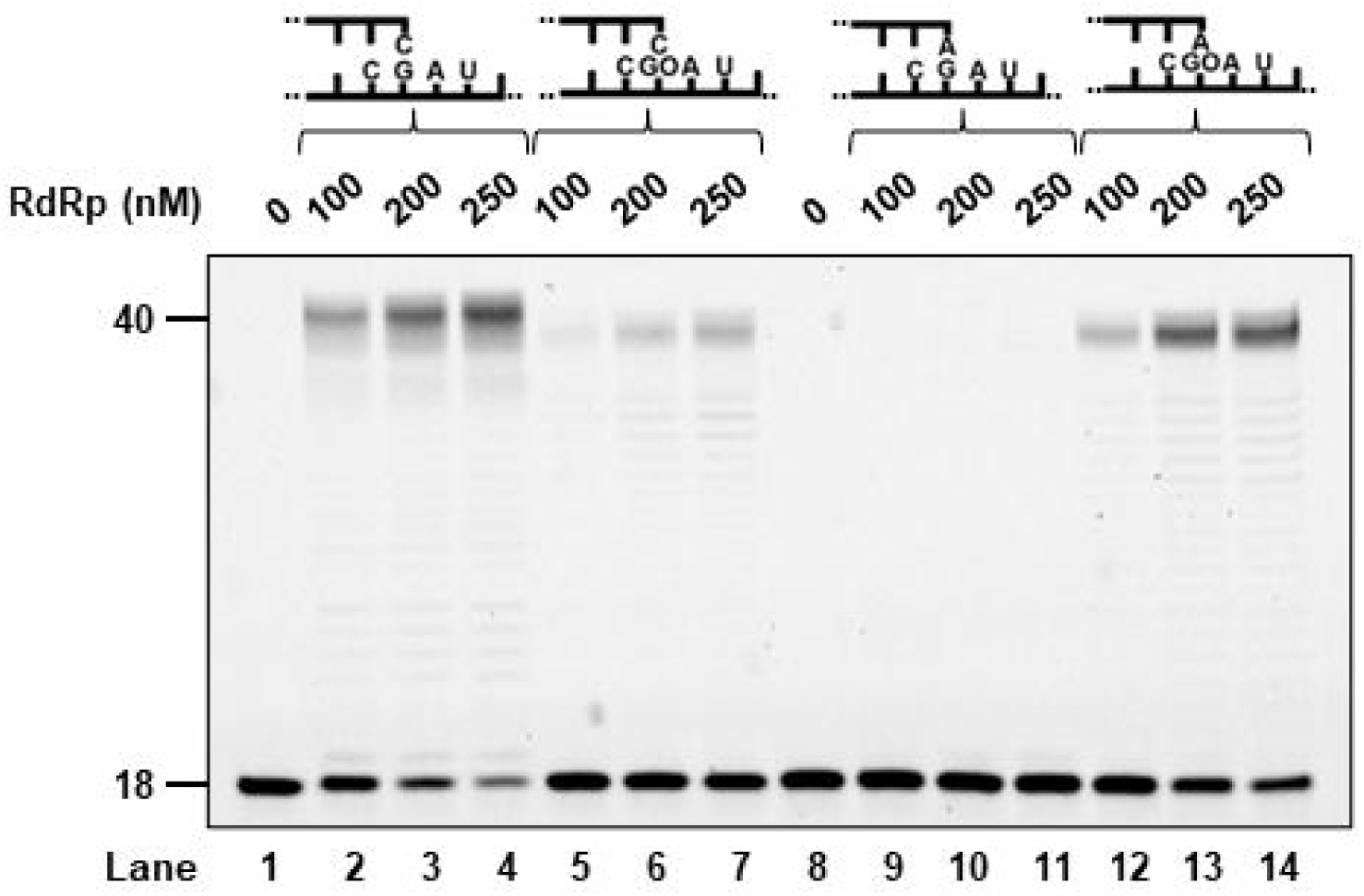
The efficiency of RdRp-mediated primer extension from the rA:8-oxo-rG termini was greater than that from rC:8-oxo-rG termini. Primer extension reactions were performed using rC:rG (lane 2-5), rA:rG (lane 6-9), rC:8-oxo-rG (GO) (lane 11–14), or rA:8-oxo-rG (GO) (lane 15–18) primer termini, all catalyzed by RdRp. Reactions were conducted with varying concentrations of RdRp (100, 200, 250, or 500 nM) for 30 min in the presence of 50 µM of the four rNTPs.

#### Standard primer extension assay

Reactions (10 µL) were conducted in a buffer consisting of 20 mM HEPES (pH 7.5), 15 mM KCl, 2 mM MgCl_2_, 1 mM DTT, and 5 % glycerol. Reaction mixtures containing 100 nM P/T RNA and 50 µM rNTPs, or each individual rNTP (rATP, rCTP, rGTP, or rUTP), were preincubated for 5 min at 37°C. Following preincubation, the primer extension reaction was initiated by adding RdRp to the reaction mixture. After 30 min, the reactions were halted by the addition of an equal amount of formamide dye, which contained 95% formamide, blue dextran (25 mg/ml), and EDTA (10 mM). The products were then resolved by electrophoresis on a 15% denaturing polyacrylamide gel (30 × 40 × 0.05 cm) for 2 h at 1,500 V. Images were captured using the iBright FL1500 Imaging Systems (Thermo Fisher Scientific).

#### Determination of the multiple turnover rate of RdRp

A 5′-Cy3-labeled 14-mer primer and a 40-mer template were utilized for the reaction. The P/T RNA at a concentration of 100 nM was equilibrated at 37°C in a buffer consisting of 20 mM HEPES (pH 7.5), 15 mM KCl, 2 mM MgCl_2_, 1 mM DTT, 5% glycerol, and 100 µM of each of the four rNTPs. Reactions were initiated by adding 100 nM RdRp. At various time intervals, aliquots were withdrawn and mixed with an equal amount of formamide dye containing blue dextran (25 mg/ml) and EDTA (10 mM). The resulting products were resolved by electrophoresis on a 15% denaturing polyacrylamide gel and analyzed as described above. The time courses of the extension reactions were determined by fitting the data to a model incorporating rising exponential and linear terms (Equation 1), which provides the first-order rate constant (*k*_obs_), the amplitude of the burst (A_0_, y-intercept), and the slope of the linear steady-state phase (*v*_ss_), as previously described (56). The multiple turnover rate (*k*_off_) of RdRp was calculated by dividing *v*_ss_ by A_0_.

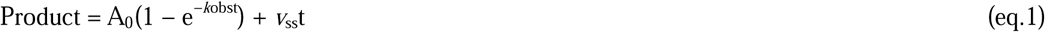

#### Michaelis–Menten steady-state kinetic analyses

The steady-state kinetic parameters for the incorporation of rG and 8-oxo-rG, as well as the extension from the 3′-terminus, were determined at 37°C using varying concentrations of individual rNTPs, as described previously (38). To assess the steady-state rate, RdRp concentration and reaction time were selected to ensure that substrate depletion or product inhibition did not affect the observed rate. The products were resolved by electrophoresis on a 20% polyacrylamide gel for 3 h at 1,700 V and analyzed as previously described. The rate of nucleotide insertion was plotted against rNTP concentrations, and the Michaelis– Menten constants (*K*_m_) and *V*_max_ values were calculated by fitting the rate data to the Michaelis–Menten equation. The *k*_cat_ value was derived by dividing the *V*_max_ values by the enzyme concentrations.

## Results

### 8-Oxo-riboguanine strongly retards the RdRp-mediated primer extension

To examine the RdRp activity of SARS-CoV-2 *in vitro*, the protein complex composed of the nsp7, nsp8, and nsp12 subunits was expressed in *E. coli* and purified through three chromatography steps: cobalt affinity, anion exchange, and size exclusion columns. The SDS–PAGE results displayed three bands corresponding to the theoretical molecular masses of nsp7 (9.37 kDa), nsp8 (22.4 kDa), and nsp12 (107 kDa), respectively (Fig 1A), indicating that the RdRp complex was purified to near homogeneity. To verify the activity of the purified RdRp complex, primer extension reactions were performed in the presence of all four rNTPs and varying amounts of RdRp, using a 40-mer template annealed to its complementary 14-mer primer, which was labeled with a cy3 fluorophore at the 5′-terminus. With the unmodified RNA template, RdRp extended the primers to form fully extended products in a concentration-dependent manner (Fig 1B, lane 1–5). The formation of fully elongated bands with minimal intermediate-length products indicates processive RNA synthesis catalyzed by RdRp. When an 8-oxo-rG-modified RNA template was used in the reactions, the primer extension reaction catalyzed by RdRp was strongly blocked by one base before the 8-oxo-rG (Fig 1B, lane 6–10).

To investigate the behavior of RdRp when encountering an oxidative lesion during primer extension, we evaluated whether RdRp remained bound at the 8-oxo-rG site on the template RNA or switched to another primer/template RNA relative to reinitiate RNA synthesis. Under conditions with an excess of rNTP and primer/template RNA relative to RdRp, the time courses of product formation displayed a biphasic pattern: an initial exponential phase followed by a linear steady-state phase. The rate of product release (*k*_off_) can be determined from the slope of the steady-state phase. The primer extension reaction was conducted over varying time intervals (Fig 1C, D). As expected, with the unmodified RNA template, quantification of fully extended products revealed a fast initial phase of product formation, followed by a linear steady-state phase (Fig 1E). Based on this linear steady-state phase, the *k*_off_ was determined to be 4.9 × 10^−3^/min (Table 1), indicating a slow rate of product release after elongation to the end of the template. In the presence of 8-oxo-rG at the 18th position of the RNA template, RdRp was strongly blocked one base before the 8-oxo-rG (Fig 1F). Based on the quantification of 17-mer products relative to the total primer amount, the *k*_off_ value of RdRp stalled before 8-oxo-rG was determined to be 2.4 × 10^−3^/min (Table 1). These results indicate that the turnover rate was significantly slower when replication was blocked by 8-oxo-rG, similar to which was observed when elongation reached the template terminus.

**Table 1.** Multiple turnover determination of RNA synthesis catalyzed by RdRp.

| | $k_{\text{obs}}$ ($\text{min}^{-1}$) | $v_{\text{ss}}$ ($\text{nM} \cdot \text{min}^{-1}$) | $k_{\text{off}}$ ($\text{min}^{-1}$) |
| --- | --- | --- | --- |
| G | $1.8 \pm 0.28$ | $0.49 \pm 0.086$ | $(4.9 \pm 0.86) \times 10^{-3}$ |
| 8-oxo-rG | $1.1 \pm 0.18$ | $0.24 \pm 0.091$ | $(2.4 \pm 0.91) \times 10^{-3}$ |

The first order rate constant ( $k_{\text{obs}}$ ) and the slope of the linear steady-state phase ( $v_{\text{ss}}$ ) were determined as described in Materials and Methods. The multiple turnover rate ( $k_{\text{off}}$ ) of RdRp was calculated by dividing $v_{\text{ss}}$ by the amplitude of the burst ( $A_0$ ).

### RdRp incorporates rCMP with a lesser extent of rAMP opposite 8-oxo-rG

It was notable that substantial amount of 18-mer and fully extended products were observed during primer extension on the template containing an 8-oxo-rG (Fig 1B, lane 6-10), indicating that RdRp was able to incorporate rNMP opposite 8-oxo-rG to some extent. Therefore, we wished to determine the specificity for the incorporation of rNMP opposite 8-oxo-rG compared to rG catalyzed by RdRp. With the unmodified RNA template, rCMP was preferentially incorporated followed by rUMP opposite rG, and the incorporation of rAMP opposite rG was not detectable (Fig 2, lane 2-5). Using an 8-oxo-rG-modified RNA template, RdRp incorporated rCMP and a lesser amount of rAMP opposite 8-oxo-rG (Fig 2, lane 6-9). The steady-state kinetic analysis was performed to more accurately measure the frequency of rNMP incorporation (*F*_ins_) opposite rG and 8-oxo-rG. The *F*_ins_ for rCMP incorporation opposite rG (1.0) was five orders of magnitude higher than that for rAMP incorporation (2.7 × 10^-6^) (Table 2). In contrast, the *F*_ins_ for rCMP incorporation opposite 8-oxo-rG (2.2 × 10^-5^) was only ∼6.5-fold higher compared to that for rAMP (3.4 × 10^-6^) (Table 2). Collectively, the frequency of rCMP incorporation opposite 8-oxo-rG was ∼4.5×10^4^-fold lower than that opposite rG. On the other hand, the frequency of rAMP incorporation opposite 8-oxo-rG was comparable to that opposite rG.

**Table 2.** Kinetic parameters for rCTP or rATP insertion opposite rG or 8-oxo-rG catalyzed by RdRp.

| rNTP:X | $K_{\text{m}}$ ($\mu\text{M}$) | $k_{\text{cat}}$ ($\text{min}^{-1}$) | $k_{\text{cat}} / K_{\text{m}}$ ($\text{min}^{-1} \cdot \mu\text{M}^{-1}$) | $F_{\text{ins}}$ |
| --- | --- | --- | --- | --- |
| rCTP:rG | $0.0021 \pm 0.00019$ | $0.020 \pm 0.0023$ | $9.6 \pm 1.1$ | 1.0 |
| rATP:rG | $100 \pm 34$ | $0.0027 \pm 0.00026$ | $(2.6 \pm 0.25) \times 10^{-5}$ | $2.7 \times 10^{-6}$ |
| rCTP:8-oxo-rG | $46 \pm 3.7$ | $0.0096 \pm 0.00052$ | $(2.1 \pm 0.11) \times 10^{-4}$ | $2.2 \times 10^{-5}$ |
| rATP:8-oxo-rG | $140 \pm 20$ | $0.0044 \pm 0.00069$ | $(3.2 \pm 0.51) \times 10^{-5}$ | $3.4 \times 10^{-6}$ |

The Michaelis-Menten constants ( $K_{\text{m}}$ ) and $V_{\text{max}}$ values were determined by fitting the data to the Michaelis-Menten equation. The $k_{\text{cat}}$ value was calculated by dividing the $V_{\text{max}}$ values by the enzyme concentrations. Frequency of rNMP insertion ( $F_{\text{ins}}$ ) was estimated by the following equation: $F_{\text{ins}} = (k_{\text{cat}}/K_{\text{m}})/(k_{\text{cat}}/K_{\text{m}})_{\text{correct pair = rC:rG}}$ . Data were expressed as mean ± S.E. obtained from three independent experiments.

### Proficient primer extension from rA:8-oxo-rG termini catalyzed by RdRp

To further investigate the ability of RdRp to bypass 8-oxo-rG, we examined the efficiency of extension from the inserted nucleotide (rCMP or rAMP) opposite 8-oxo-rG in the presence of four rNTPs. Using the unmodified RNA template, RdRp efficiently extended from the matched (rC:rG) termini (Fig 3, lane 2-4), while almost no extension occurred from the mismatched (rA:rG) termini (Fig 3, lane 9-11). Notably, when an 8-oxo-rG-modified RNA template was used, RdRp showed greater activity in extending from the rA:8-oxo-rG termini (Fig 3, lane 5-7 and 12-14). Quantitative analysis through steady-state kinetic measurements revealed that the frequency of extension from the primer termini (*F*_ext_) for rA:rG (1.7 × 10^−6^) was five orders of magnitude lower than that for rC:rG (1.0) (Table 3). In contrast, the *F*_ext_ value for rA:8-oxo-rG (1.4 × 10^−3^) was approximately 45-fold higher than that for rC:8-oxo-rG (3.2 × 10^−5^). By combining the steady-state kinetic parameters for nucleotide insertion (*F*_ins_) and chain extension (*F*_ext_), the relative bypass frequency (*F*_ins_ × *F*_ext_) past 8-oxo-rG was determined. The *F*_ins_ × *F*_ext_ value for rC:rG (1.2 × 10^2^) was eleven orders of magnitude higher than that for rA:rG (5.4 × 10^−10^) (Table 4). However, the *F*_ins_ × *F*_ext_ for rC:8-oxo-rG (8.2 × 10^−8^) was approximately seven-fold lower than that for rA:8-oxo-rG (5.7 × 10^−7^) (Table 4).

**Table 3.** Kinetic parameters for extension from the primer termini catalyzed by RdRp*^a^*.

| N:X | $K_m$ (μM) | $k_{cat}$ ($\text{min}^{-1}$) | $k_{cat} / K_m$ ($\text{min}^{-1} \cdot \mu\text{M}^{-1}$) | $F_{ext}$ |
| --- | --- | --- | --- | --- |
| rC/rG | $0.013 \pm 0.0043$ | $0.16 \pm 0.016$ | $12 \pm 1.2$ | 1.0 |
| rA/rG | $250 \pm 240$ | $0.0025 \pm 0.00057$ | $(2.1 \pm 0.23) \times 10^{-5}$ | $1.7 \times 10^{-6}$ |
| rC/8-oxo-rG | $97 \pm 18$ | $0.038 \pm 0.0034$ | $(3.9 \pm 0.36) \times 10^{-4}$ | $3.2 \times 10^{-5}$ |
| rA/8-oxo-rG | $2.3 \pm 0.59$ | $0.041 \pm 0.0051$ | $(1.8 \pm 0.22) \times 10^{-2}$ | $1.4 \times 10^{-3}$ |

<sup>a</sup> Steady-state kinetic parameters were determined for incorporation of rUMP opposite template adenosine (rA) adjacent to rC/rG, rA/rG, rC/8-oxo-rG, or rA/8-oxo-rG primer termini. The Michaelis-Menten constants ( $K_m$ ) and $V_{max}$ values were determined by fitting the rate data to the Michaelis-Menten equation. The $k_{cat}$ value was calculated by dividing the $V_{max}$ values by the enzyme concentrations. Frequency of extension from the primer termini ( $F_{ext}$ ) was estimated by the following equation: $F_{ext} = (k_{cat}/K_m)/(k_{cat}/K_m)[\text{correct pair} = \text{rC:rG}]$ . Data were expressed as mean ± S.E. obtained from three independent experiments.

**Table 4.** Relative bypass frequency catalyzed by RpRp.

| N:X | $F_{ins} \times F_{ext}$ |
| --- | --- |
| rC:rG | 1.0 |
| rA:rG | $4.6 \times 10^{-12}$ |
| rC:8-oxo-rG | $7.0 \times 10^{-10}$ |
| rA:8-oxo-rG | $4.8 \times 10^{-9}$ |

## Discussion

ROS was generated in host cells in response to viral infection (24–28), making viral genomic RNA susceptible to oxidation in infected cells. G to U transversion mutations have frequently been observed in the SARS-CoV-2 genome (13), which may be linked, at least in part, to RdRp-catalyzed replication past 8-oxo-rG. To gain a deeper understanding of viral RNA mutagenesis, it is essential to clarify the potential role of RdRp in translesion synthesis across RNA lesions.

In primer extension reactions using the purified RdRp complex and primer/template RNA containing a single 8-oxo-rG, RdRp-mediated RNA synthesis was significantly hindered, stalling one base before the 8-oxo-rG (Fig 1B, lane 6–10). Quantitative analyses showed that the efficiency of bypassing 8-oxo-rG was approximately eight orders of magnitude lower than that of undamaged rG (Table 4). These findings suggest that ribonucleotide oxidation results in a replication-blocking lesion in the viral RNA genome. Similarly, studies have shown that 8-oxo-rG embedded in DNA strongly inhibits DNA synthesis catalyzed by human pol α and pol κ (38), whereas oxidative deoxyguanosine, (8-oxo-dG) is readily bypassed by these pols (38, 57–59). These observations suggest that the different sugar backbones of 8-oxo-rG and 8-oxo-dG influence the structural and dynamic behavior of oxidative base lesions. In RNA, the ribonucleotide adopts a C3′-endo sugar pucker conformation, whereas in DNA, the deoxyribonucleotide adopts a C2′-endo conformation (60). Since sugar pucker conformation affects the positioning of the base (60), it is likely that RNA lesions have differential miscoding properties compared to their DNA counterparts. When 8-oxo-dG adopts an *anti-* conformation and forms Watson–Crick pairs with cytosine in DNA, a steric clash may occur between the C8-oxygen of 8-oxo-dG and the O4′ of deoxyribose (33). To avoid this steric hindrance, 8-oxo-dG shifts to *a syn*-conformation, forming stable Hoogsteen base pairs with adenine, which leads to G:C to T:A transversion mutations. In contrast, during RdRp replication past 8-oxo-rG, steady-state kinetic analysis revealed that the efficiency of rAMP insertion opposite the lesion (3.2 × 10^−5^) was similar to that opposite rG (2.6 × 10^−5^) (Table 2). These results suggest that the ribose backbone of 8-oxo-rG in the template hinders stable 8-oxoG:A base pairing within the catalytic site. Structural analysis of the SARS-CoV-2 RdRp complex (61, 62), reveals that residues N496, K500, and N577 in the finger domain, along with Y595 in the palm domain, play a role in stabilizing the phosphate backbone of the template prior to rNMP insertion. The template base is further stabilized by interactions with the backbone of S682 and the K545 side chain. The reduced efficiency of rAMP incorporation opposite 8-oxo-rG suggests that the accommodation of 8-oxo-rG in the *syn*-conformation may cause steric hindrance with the side chains of active site residues during nucleotide insertion, thereby preventing the formation of a stable Hoogsteen base pair.

While RdRp preferentially incorporated rCMP over rAMP opposite 8-oxo-rG (Fig 2, Table 2), the efficiency of extension from the rA:8-oxo-rG termini was significantly higher than from the rC:8-oxo-rG termini (Fig 3, Table 3). Considering both the frequencies of nucleotide insertion opposite 8-oxo-rG and chain extension from the 3′-primer termini, the *F*_ins_ × *F*_ext_ for rA:8-oxo-rG was seven-fold higher than for rC:8-oxo-rG (Table 4). This suggests that RdRp can bypass 8-oxo-rG in an error-prone manner, potentially leading to G to U transversion mutations in the viral genome. Based on our findings, Figure 4 proposes a model of RNA replication by RdRp across oxidative RNA damage. An 8-oxo-rG lesion in the template RNA strongly impedes RNA synthesis, though RdRp can partially bypass it. RdRp preferentially incorporates rCMP, followed by rAMP, opposite 8-oxo-rG. However, the efficiency of extension from rC:8-oxo-rG termini is lower than that from rA:8-oxo-rG, indicating that 8-oxo-rG acts as both a replication-blocking RNA lesion and a promutagenic base that can cause errors during viral genome replication. Given that the catalytic domain of RdRp is highly conserved across RNA viruses despite sequence differences (63), it is likely that 8-oxo-rG induces similar miscoding effects in RdRp from various RNA viruses. Our kinetic analyses show that the relative bypass frequency of RdRp past 8-oxo-rG is extremely low compared to undamaged rG (Table 4). To counteract the detrimental effects of oxidative RNA lesions, other viral and cellular replisome components may assist in RNA replication progression. SARS-CoV-2 RdRp has been reported to interact with additional nsps such as the proofreading exonuclease nsp14, its activator nsp10, and the single-stranded nucleic acid-binding protein nsp9, forming a replication–transcription complex (22, 64, 65). RdRp also associates with host-derived proteins, including HSC20 and components involved in *de novo* Fe–S cluster assembly and biogenesis (66). These factors may influence the efficiency and fidelity of RdRp-mediated translesion RNA synthesis. Further investigation is needed to fully understand the mechanisms underlying RNA damage tolerance in viruses.

**Fig 4.**
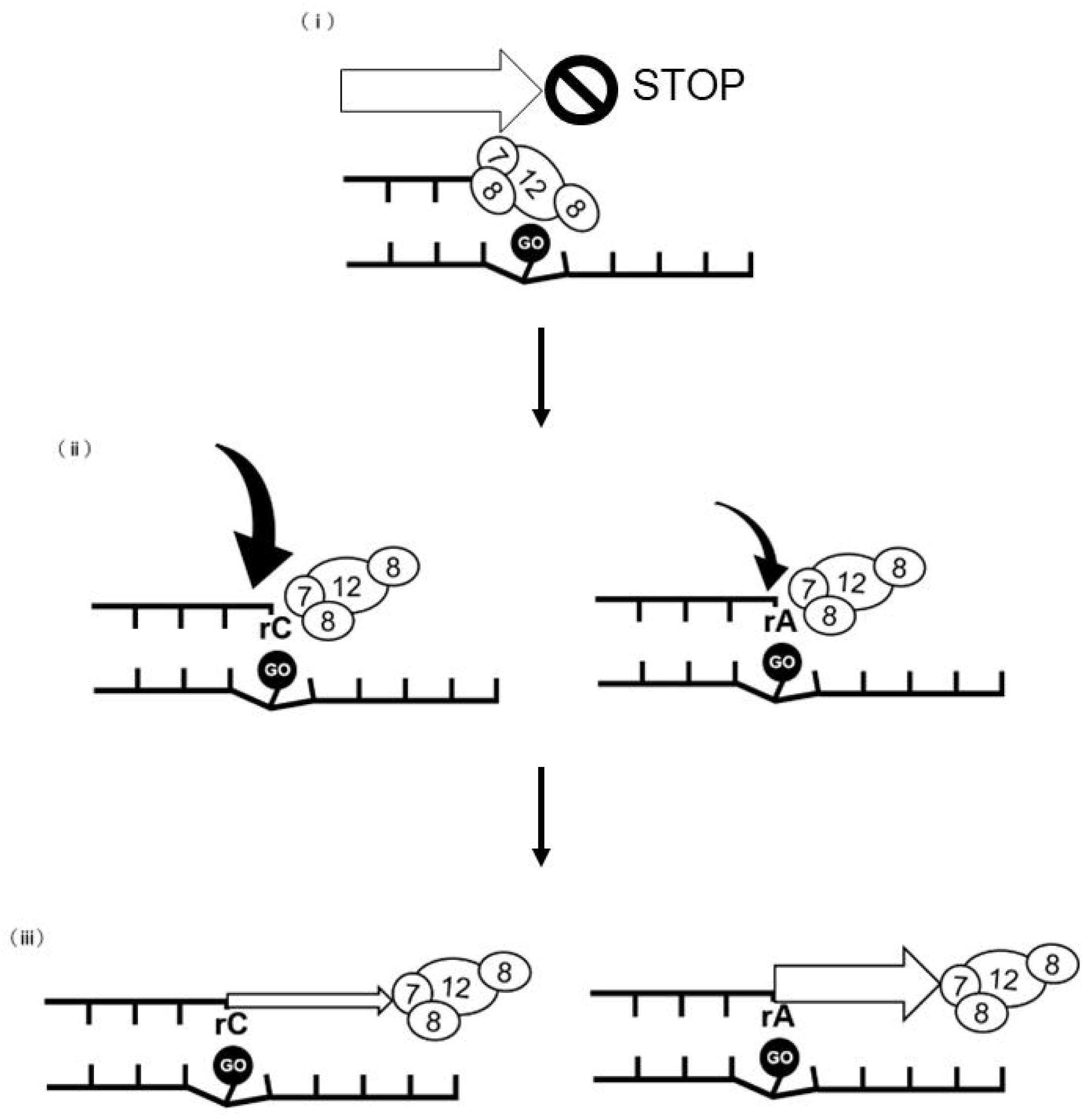
Scheme for RNA synthesis across the oxidative RNA lesion catalyzed by RdRp. (L) An 8-oxo-rG (GO) on the template RNA significantly inhibits RdRp-mediated RNA synthesis. (L) Once RdRp bypasses 8-oxo-rG, it preferentially incorporates rCMP, with a smaller amount of rAMP incorporated opposite the lesion. (L) RdRp demonstrates greater activity when extending from the mutagenic rA:8-oxo-rG pair compared to the lower efficiency of extension from the non-mutagenic rC:8-oxo-rG termini. Table 1. Multiple turnover determination of RNA synthesis catalyzed by RdRp.

## Abbreviations

(RdRp): RNA-dependent RNA polymerase
ROS: (Reactive oxygen species)
HCV: (Hepatitis C virus)
HIV: (Human immunodeficiency virus)
NiRAN: (Nidovirus RdRp-associated nucleotidyltransferase), 7,8-dihydro-8-oxo-riboguanosine (8-oxo-rG)
(CoVs): coronaviruses
(SARS-CoV-2): severe acute respiratory syndrome coronavirus 2

## Data availability

All data are contained within the manuscript.

## Acknowledgments

We are grateful to Dr. Masayuki Yokoi (Kobe University) for his valuable suggestions and to Dr. Petr Grúz (National Institute of Health Sciences) for his technical support.

## Author contributions

M.A. and A.S. designed the research; M.A. and A.S. established the experimental setup; M.A., K.S., K.U., and A.S. contributed to discussions about the study; M.A. and A.S. conducted the research and analyzed the data; M.A. and A.S. wrote the manuscript; K.S., K.U., and A.S. provided research direction; K.S. and K.U. offered feedback on the results and manuscript throughout the process; M.A., K.S., K.U., and A.S. revised the paper; all authors approved the final version of the manuscript.

